# What do adversarial images tell us about human vision?

**DOI:** 10.1101/2020.02.25.964361

**Authors:** Marin Dujmović, Gaurav Malhotra, Jeffrey Bowers

## Abstract

Deep convolutional neural networks (DCNNs) are frequently described as promising models of human and primate vision. An obvious challenge to this claim is the existence of *adversarial images* that fool DCNNs but are uninterpretable to humans. However, recent research has suggested that there may be similarities in how humans and DCNNs interpret these seemingly nonsense images. In this study, we reanalysed data from a high-profile paper and conducted four experiments controlling for different ways in which these images can be generated and selected. We show that agreement between humans and DCNNs is much weaker and more variable than previously reported, and that the weak agreement is contingent on the choice of adversarial images and the design of the experiment. Indeed, it is easy to generate images with no agreement. We conclude that adversarial images still challenge the claim that DCNNs constitute promising models of human and primate vision.

## 1. Introduction

Deep convolutional neural networks (DCNNs) are models of computer vision that have reached, and in some cases exceeded, human performance in many image classification benchmarks such as `ImageNet` [18]. In addition to having obvious commercial implications, these successes raise questions as to whether DCNNs identify objects in a similar way to the infereotemproal cortex (IT) that supports object recognition in humans and primates. If so, these models may provide important new insights into the underlying computations performed in IT. Consistent with this possibility, a number of researchers have highlighted various functional similarities between DCNNs and human vision [29] as well as similarities in patterns of activation of neurons in IT and units in DCNNs [35]. This has led some authors to make strong claims regarding the theoretical significance of DCNNs to neuroscience and psychology. For example, Kubilius et al. [24] write: “Deep artificial neural networks with spatially repeated processing (a.k.a., deep convolutional [Artificial Neural Networks]) have been established as the best class of candidate models of visual processing in primate ventral visual processing stream” (p.1).

One obvious problem in making this link is the existence of *adversarial* images which are designed to fool DCNNs. Figure 1 shows examples of two types of adversarial images. On first impression, it seems inconceivable that these adversarial images would ever confuse humans. There is now a small industry of researchers creating adversarial attacks that produce images which DCNNs classify in bizarre ways [1]. The confident classification of these adversarial images by DCNNs suggests that humans and current architectures of DCNNs perform image classification in fundamentally different ways. If this is the case, the existence of adversarial images poses a challenge to research that considers DCNNs as models of human behaviour [e.g., 24, 33, 28, 11, 13], or as plausible models of neural firing patterns in primate and human visual cortex [e.g., 22, 10, 31, 36, 14, 12].

**Figure 1:**
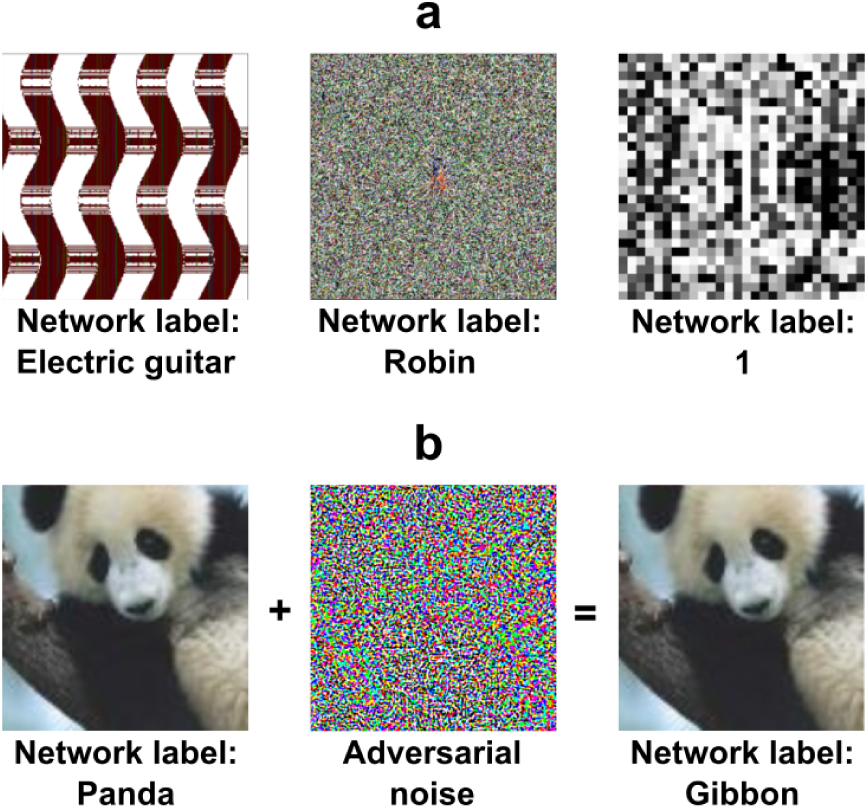
Examples of two types of adversarial images. (a) *fooling adversarial images* taken from [26] that do not look like any familiar object. The two images on the left (labelled ‘Electric guitar’ and ‘Robin’) have been generated using evolutionary algorithms using *indirect* and *direct* encoding, respectively, and classified confidently by a DCNN trained on `ImageNet`. The image on the right (labelled ‘1’) is also generated using an evolutionary algorithm using *direct* encoding and it is classified confidently by a DCNN trained on `MNIST`. (b) An example of a *naturalistic adversarial image* taken from [17] that is generated by perturbing a naturalistic image on the left (classified as ‘Panda’) with a high-frequency noise mask (middle) and confidently (mis)classified by a DCNN (as a ‘Gibbon’).

However, some recent studies have suggested that there may, in fact, be a larger overlap between DCNNs and humans in how they process these adversarial images. Zhou and Firestone [37] (*Z&F* from here on) recently reported that humans can reliably *decipher* fooling adversarial images that, on first viewing, look uninterpretable (as in Figure 1a). The authors took a range of published adversarial images that were claimed to be uninterpretable by humans and, in a series of experiments, they reported that a high percentage of participants (often close to 90%) classified the images in the same way as the DCNNs at an above chance level.

Here we show these conclusions are not justified. We first identified problems with the *Z&F* analyses, and when corrected, the agreement between humans and DCNNs was much weaker and highly variable between participants and images. The remaining agreement appeared to reflect participants making educated guesses based on some superficial features (such as colour) within images and the limited response alternatives presented to them. We then carried out four experiments in which we systematically manipulated factors that can contribute to an observed agreement between humans and DCNNs in order to better understand the how humans interpret adversarial images. The experiments demonstrate that the overlap between human and DCNN classification is contingent upon various details of the experimental design such as the selection of adversarial images used as stimuli, the response alternatives presented to participants during the experiment, the adversarial algorithm used to generate the images and the dataset on which the model was trained. When we controlled for these factors, we observed that the agreement between humans and DCNNs dropped to near chance (or even below chance) levels. We also show that it is straightforward to generate adversarial images that fool networks trained on `ImageNet` but are truly meaningless to human participants, irrespective of how the stimuli are selected or response alternatives are presented to a participant. We take the findings to highlight a dramatic difference between human and DCNN object recognition.

## 2. Results

### 2.1. Reassessing the level of agreement in Zhou and Firestone [37]

Our first step, in trying to understand the surprisingly large agreement between humans and DCNNs observed by *Z&F*, was to reassess how they measured this agreement on classification of adversarial images. *Z&F* conducted seven experiments in which they measured agreement by computing the number of trials on which the participants matched the DCNN’s classification and working out whether this number is numerically above or below chance level. So in Experiment 3, for example, a participant is shown an adversarial image on each trial and asked to choose one amongst 48 labels for that image. Chance level is 1/48, so if the participant chooses the same label as the DCNN on two or more trials, they were labelled as agreeing with the DCNN. In addition, half of the participants who agreed with the DCNN on only 1/48 trials were also counted towards the number of participants who agreed with the DCNN. When computed in this manner, *Z&F* calculated that 142 out of 161 (88%) participants in Experiment 3a, and 156 out of 174 (90%) in Experiment 3b agreed with the DCNN at above chance levels.

But these rates of agreement are both misleading and statistically unsound as they ignore inter-individual variability and assign the same importance to a participant that agrees on 2 out of 48 trials as a participant who agrees on all 48 trials with the DCNN. In fact, not a single participant in Experiments 3a&b (from a total of 335 participants) agreed with the model on a majority (24 or more) of trials, yet the reported level of agreement is nearly 90%. In fact, if participants were blindfolded and responded randomly on each of 48 trials, it can be shown that this method of analysis will result in a ∼ 45% agreement between the DCNN and these blindfolded participants who have never seen the stimuli (see Methods). Using a statistic that treats 45% agreement as chance is liable to be misinterpreted by readers.

A more appropriate statistical measure would be to establish whether a participant’s choices are *significantly* different from a chance agreement with the DCNN – that is, carry out a binomial test on each participant’s data. Measured in this manner (independent 2-tailed binomial tests with a critical p-value of 0.05 for each participant), the agreement in Experiment 3a between DCNN and participants drops from 88% to 57.76%. In Table 1, we summarise the results of carrying out the more appropriate binomial test for all seven experiments when assessing agreement over participants and images. (Note, this is different from the binomial test carried out by *Z&F*, who first label participants as agreeing with DCNN in the manner described above and then used the binomial test to measure whether the proportion of such participants is above chance).

**Table 1:** Different ways of measuring agreement between humans and DCNNs in experiments conducted by Zhou and Firestone [37]).

| Exp. | Test type | Participants |  | Images |  |
| --- | --- | --- | --- | --- | --- |
|  |  | ZF19 | Binomial | ZF19 | Binomial |
| 1 | Fooling 2AFC <sup>N15</sup> | 98% | 85.4% | 94% | 85.42% |
| 2 | Fooling 2AFC <sup>N15</sup> | 91% | 34.25% | 71% | 56.25% |
| 3a | Fooling 48AFC <sup>N15</sup> | 88% | 57.76% | 79% | 56.25% |
| 3b | Fooling 48AFC <sup>N15</sup> | 90% | 63.22% | 81% | 56.25% |
| 4 | TV-static 8AFC <sup>N15</sup> | 81% | N/A | 100% | 75% |
| 5 | Digits 9AFC <sup>P16</sup> | 89% | 34.75% | 73% | 33% |
| 6 | Naturalistic 2AFC <sup>K18</sup> | 87% | N/A | 100% | 81.82% |
| 7 | 3D Objects 2AFC <sup>A17</sup> | 83% | 26.85% | 78% | 52.83% |

<sup>a</sup> ZF19 refers to agreement as calculated by $Z\mathcal{E}F$ while Binomial refers to percentages of participants and images for which agreement was calculated as statistically significantly above chance (2-tailed Binomial test with a critical p value of 0.05). The ‘Test type’ refers to the type of adversarial images and number of response alternatives presented in the experiment by $Z\mathcal{E}F$ .
<sup>b</sup> The number of images in Experiment 4 and 6 were too low (eight and ten) for it to make sense to conduct a statistical test (thus the N/A value).
<sup>c</sup> Stimuli sources: N15 - Nguyen et al. [26]; P16 - Papernot et al. [27]; K18 - Karmon et al. [21]; A17 - Athalye et al. [3]

Another way of measuring agreement is to simply report the average agreement. This can be calculated as the mean percentage of images (across participants) on which participants and DCNNs agree. A key advantage of this measure is that it takes into consideration the level of agreement of each participant (a participant who agrees on 4/48 trials is not treated equivalently to a participant who agrees on 48/48 trials). *Z&F* only reported mean agreement for the first of their seven experiments and in Table 2 we report mean agreement levels in all experiments. In all cases the mean agreement levels indicate much more modest agreement.

**Table 2:** Mean agreement in the experiments conducted by Zhou and Firestone [37])

| Exp. | Mean agreement | Chance |
| --- | --- | --- |
| 1 | 74.18% (35.61 out of 48 images) | 50% |
| 2 | 61.59% (29.56 out of 48 images) | 50% |
| 3a | 10.12% (4.86 out of 48 images) | 2.08% |
| 3b | 9.96% (4.78 out of 48 images) | 2.08% |
| 4 | 28.97% (2.32 out of 8 images) | 12.5% |
| 5 | 16% (1.44 out of 9 images) | 11.11% |
| 6 | 73.49% (7.3 out of 10 images) | 50% |
| 7 | 59.55% (31.56 out of 53 images) | 50% |

To give the readers a sense of the levels of agreement observed in these experiments, we have also computed the average number of images in each experiment where humans and DCNNs agree as well as the level of agreement expected if participants were responding at chance.

### 2.2. Reassessing the basis of the the agreement in Zhou and Firestone [37]

While both binomial test (Table1) and mean agreement (Table 2) highlight a more modest level of agreement, it is nevertheless the case that even these methods show that the overall agreement was above chance. Perhaps the most striking result is in *Z&F*’s Experiment 3 where participants had to choose between 48 response alternatives and mean agreement was ∼ 10% with chance being ∼ 2%.

In order to clarify the basis of overall agreement we first assessed the level of agreement for each of the 48 images separately. As shown in Supplementary Figure 1.1, the distribution of agreement levels was highly skewed and had a large variance. There was a small subset of images that looked like the target class (such as the Chainlink Fence, which can be seen in Supplementary Figure 1.2) and participants showed a high level of agreement with DCNNs on these images. Another subset of images with lower (but statistically significant) levels of agreement contained some features consistent with the target class, such as the Computer Keyboard which contains repeating rectangles. But agreement on many images (21*/*48) was at or below chance levels. This indicates that the agreement is largely driven by a subset of adversarial images, some of which (such as the Chainlink Fence) simply depict the target class.

We also observed that there was only a small subset of images on which participants showed a clear preference amongst response alternatives that matched the DCNN’s label. For most adversarial images, the distribution of participant responses across response alternatives was fairly flat (see Supplementary Figure 1.3) and the most frequent human response did not match the machine label even when agreement between humans and DCNNs was above chance (see Supplementary Figure 1.2). In fact, the label assigned to the image by DCNNs was ranked 9th (Experiment 3a) or 10th (Experiment 3b) on average. 75% of the adversarial images in Experiment 3a and 79.2% in Experiment 3b were not assigned the label chosen by the DCNN with highest frequency (Supplementary Figure 1.2). This indicates that most adversarial images do not contain features required by humans to uniquely identify an object category.

Collectively, these findings suggest that the above chance level of agreement was driven by two subsets of images. A very small subset of images have features that humans can perceive and are highly predictive of the target category (e.g., Chainlink Fence image that no one would call “uninterpretable”), and another subset of images that include visible features consistent with the target category as well as a number of other categories. These category-general features (such as colour or curvature) are what *Z&F* called “superficial commonalities” between images ([37], p.2). For this subset of images, the most frequent response chosen by participants does *not* usually match the label assigned by the DCNN. Participants in these cases seem to be making educated guesses using superficial features of the target images to hedge their bets. For the rest of the images agreement is at or below chance levels.

In order to more directly test how humans interpret adversarial images we carried out four experiments. First, if participants are making educated guesses based on superficial features, then agreement levels should decrease when presented with response alternatives that do not support this strategy. We test this in Experiment 1. Second, if a DCNN develops human-like representations for a subset of categories (e.g., the Chainlink Fence category for which human-DCNN agreement was high for a specific adversarial image of a chainlink fence), then it should not matter which adversarial image from these categories is used to evaluate agreement. We test this in Experiment 2. Third, if DCNNs are processing images in very different ways to humans, then it should be possible to find situations in which overall agreement levels are at absolute chance levels. In Experiment 3 we show that one class of adversarial images for the `MNIST` dataset generated overall chance level agreement. Finally, in Experiment 4 we show that it is straightforward to generate adversarial images for the `ImageNet` dataset that produce overall chance level agreement. The findings further undermine any claim that DCCNs and humans categorize adversarial images in a similar way.

### 2.3. Experiment 1: Response alternatives

One critical difference between decisions made by DCNNs and human participants in an experiment is the number of response alternatives available. For example, DCNNs trained on `ImageNet` will choose a response from amongst 1000 alternatives while participants will usually choose from a much smaller cohort. In Experiment 1, we tested whether agreement levels are contingent on how these response alternatives are chosen during an experiment. We chose a subset of ten images from the 48 that were used by *Z&F* and identified four *competitive* response alternatives (from amongst the 1000 categories in `ImageNet`) for each of these images. One of these alternatives was always the category picked by the DCNN and the remaining three were subjectively established as categories which share some superficial visual features with the target adversarial image. For example, one of the adversarial images contains a *florescent orange curve* and is confidently classified by the DCNN as a Volcano. For this image, we chose the set of response alternatives {Lighter, Missile, Table lamp, Volcano}, all of which also contain this superficial visual feature. See Supplementary Figure 2.1 for the complete list of images and response alternatives. Participants were then shown each of these ten images and asked to choose one amongst these four competitive response alternatives. Note that if humans possess a “machine-theory-of-mind”, it should not matter how one samples response alternatives as a DCNN classifies the fooling adversarial images with high confidence (*>* 99%) in the presence of *all* 999 alternative labels, including the competing alternatives we have selected. In the control condition an independent sample of participants completed the same task but the alternative labels were chosen at *random* from the 48 used by *Z&F*.

We observed that agreement levels fell nearly to chance in the competitive condition while being well above chance in the random condition (see Figure 2). The mean agreement level in the competitive condition was at 28.5% (SD = 11.67) with chance being at 25%. A single sample t-test comparing the mean agreement level to the fixed value of 25% did show the difference was significant (*t*(99) = 3.00, *p* = .0034, *d* = 0.30). However, in the random condition mean agreement was 49.8% (SD = 16.02) which was both significantly above chance (*t*(99) = 15.48, *p <* .0001, *d* = 1.54) and well above agreement in the competitive condition (*t*(198) = 10.75, *p <* .0001, *d* = 1.52). Both conditions are in stark contrast to the DCNN which classified these images with a confidence *>* 99% even in the presence of these competing categories.

**Figure 2:**
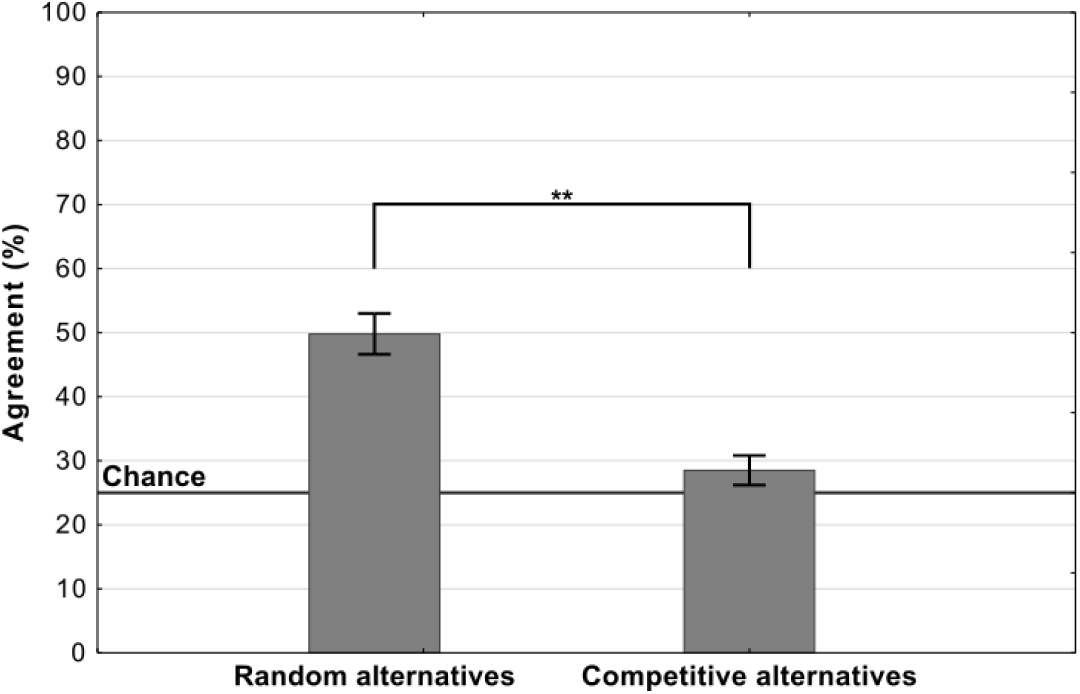
Average levels of agreement in Experiment 1 (error bars denote 95% confidence intervals).

These results highlight a key contrast between human and DCNN image classification. While the features in each of these adversarial images are sufficient for a DCNN to uniquely identify one amongst a 1000 categories, for humans they are not. Instead features within these images only allow them to identify a cohort of categories. Thus, the observed decrease in agreement between the random and competitive conditions supports the hypothesis that participants are making plausible guesses in these experiments, using superficial features (shared amongst a cohort of categories) to eliminate response alternatives.

It should be noted that *Z&F* were themselves concerned about how the choice of response alternatives may have influenced their results. Therefore, they carried out another experiment where, instead of choosing between the DCNN’s preferred label and another randomly selected label, the participants had to choose between the DCNN’s 1^*st*^ and 2^*nd*^-ranked labels. The problem with this approach is that the DCNN generally has a very high level of confidence (*>* 99%) in it’s 1^*st*^ choice. Accordingly, it is not at all clear that the 2^*nd*^ most confident choice made by the network provides the most challenging response alternative for humans. The results from Experiment 1 show that when the competing alternative is selected using a different criterion, the agreement between participants and DCNNs does indeed drop to near-chance levels.

### 2.4. Experiment 2: Target adversarial images

Our reanalysis above also showed that there was large variability in agreement between images. One possible explanation for this is that the DCNN learns to represent some categories (such as Chainlink Fence or Computer Keyboard) in a manner that closely relates to human object recognition while representations for other categories diverge. If there was meaningful overlap between human and DCNN representations for a category, we would expect participants to show a similar level of agreement on all adversarial images for this category as all adversarial images will capture these common features. So replacing an adversarial image from these categories with another image generated in the same manner should lead to little change in agreement. In Experiment 2 we directly tested this hypothesis by sampling two different images (amongst the five images for each category generated by Nguyen et al. [26]) for the same ten categories from Experiment 1. The first image was chosen such that it contained some superficial features that subjectively seemed to us to overlap the target class. We labelled these as the *best-case* images. Similarly, we selected a set of *worst-case* images which we subjectively assessed as containing the least number of superficial features overlapping with the target class. An example of each type of image is shown in Figure 3. While selecting either type of image, we only considered images that were classified by the DCNN as the respective target class with *>* 99% confidence.

**Figure 3:**
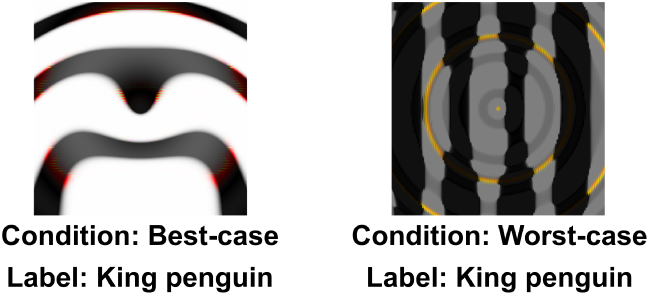
Example of *best-case* and *worst-case* images for the same category (‘penguin’) used in Experiment 2.

Figure 4 shows the mean agreement for participants viewing the *best-case* and *worst-case* adversarial images. The difference in agreement between the two conditions was highly significant (*t*(199) = 17.41, *p* < .0001, *d* = 2.46). Both groups showed agreement levels significantly different from chance (which was at 25%). The best-case group was significantly above chance (*t*(100) = 16.08, *p* < .0001, *d* = 1.60) while the worst-case was significantly below chance (*t*(99) = 7.44, *p* < .0001, *d* = 0.74).

**Figure 4:**
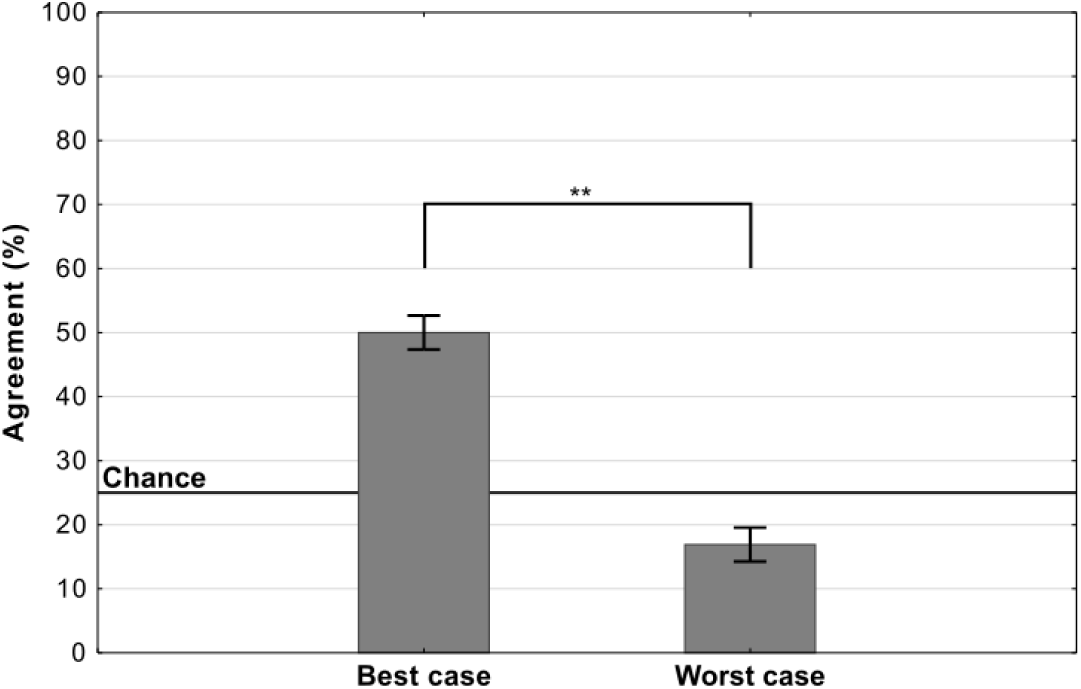
Average levels of agreement in Experiment 2 (error bars denote 95% confidence intervals).

Thus, we observed a large drop in agreement when we replaced one set of adversarial images with a different set, and there was no evidence for consistent above-chance agreement for all adversarial images from a subset of categories (see Supplementary Figure 2.2 for an item-wise breakdown). In other words, we did not observe any support for the hypothesis that DCNNs learn to represent even a subset of categories in a manner that closely relates to human object recognition.

### 2.5. Experiment 3: Different types of adversarial images

Although we can easily reduce DCNN-human agreement to chance by judicially selecting the targets and foils, it remains the case that a random selection of targets and foils has led to above chance performance on this set of images. In the next experiment, we asked whether this effect is robust across different types of adversarial images. All the images in the experiments above were generated to fool a network that had been trained on `ImageNet` and belonged to the subclass of *regular* adversarial images generated by Nguyen et al. [26] using an *indirect encoding* evolutionary algorithm. In fact, Nguyen et al. [26] generated four different types of adversarial images by manipulating the type of encoding – *direct* or *indirect* – and the type of database the network was trained on – `ImageNet` or `MNIST` (see Figure 5). We noticed that *Z&F* used images designed to fool DCNNs trained on images from `ImageNet`, but did not consider the adversarial images designed to fool a network trained on `MNIST` dataset. To our eyes, these `MNIST` adversarial images looked completely uninterpretable and we wanted to test whether the above chance agreement was contingent on which set of images were used in the experiments.

**Figure 5:**
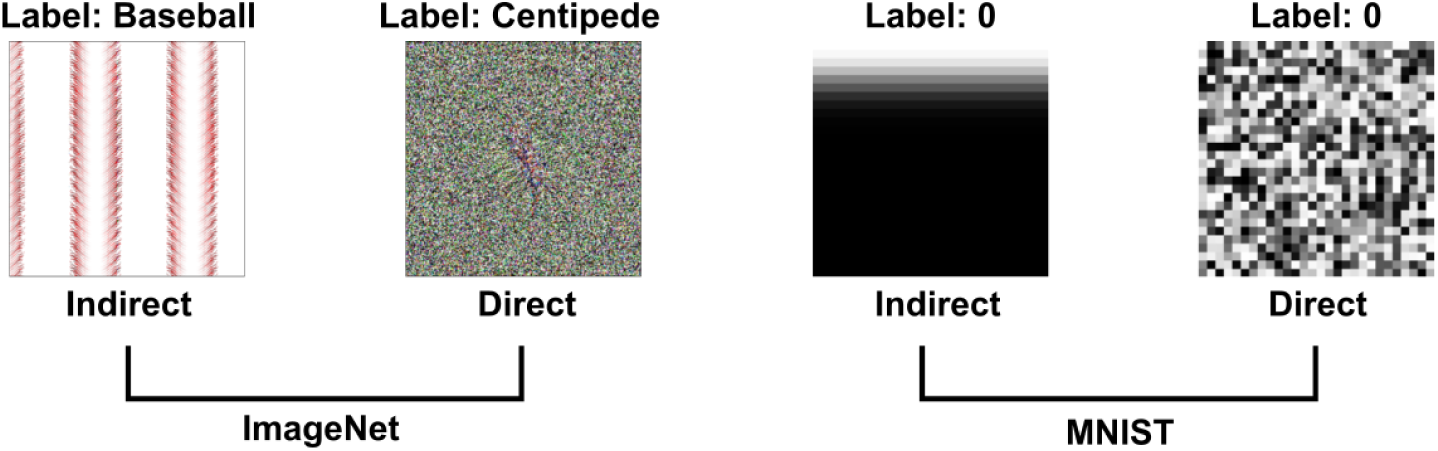
Examples of images from [26] used in the four experimental conditions in Experiment 3. Images are generated using an evolutionary algorithm either using the *direct* or *indirect* encoding and generated to fool a network trained on either `ImageNet` or `MNIST`

Accordingly, we designed a 2×2 experiment in which we tested participants on all four conditions corresponding to the four types of images (Figure 5). Since `MNIST` has ten response categories and we wanted to compare results for the `MNIST` images with `ImageNet` images, we used the same 10 categories from Experiments 1 and 2 for the two `ImageNet` conditions. On each trial, participants were shown an adversarial image and asked to choose one out of ten response alternatives that remained fixed for all trials.

Mean agreement levels in this experiment are shown in Figure 6. We observed a large difference in agreement levels depending on the types of adversarial images. Results of a two-way repeated measures ANOVA revealed a significant effect of dataset on agreement levels (*F* (1, 197) = 298.62, *p* < .0001, 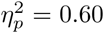). Participants agreed with DCNN classification for images designed to fool `ImageNet` classifiers significantly more than for images designed to fool `MNIST` classifiers. Participants also showed significantly larger agreement for indirectly-encoded compared to directly-encoded images (*F*(1,197) = 67.57, *p* < .0001, 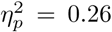). The most striking observation was that agreement dropped from 26% for `ImageNet` images to near chance for `MNIST` images. Participants were slightly above chance for indirectly-encoded `MNIST` images (*t*(197) *>* 6.30, *p <* .0001, *d* = 0.44) and at chance agreement for directly-encoded `MNIST` images (*t*(197) = 1.03, *p* = 0.31).

**Figure 6:**
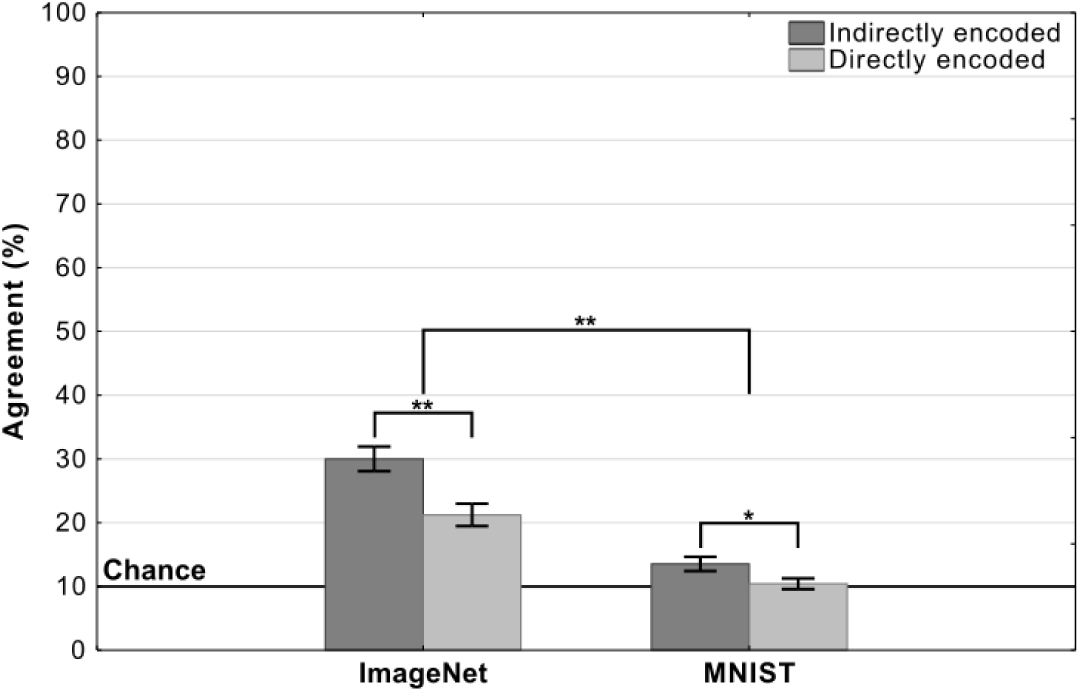
Agreement (mean percentage of images on which a participant choices agree with the DCNN) as a function of experimental condition in Experiment 3 (error bars denote 95% confidence intervals).

In addition to the between-condition differences, we also found high within-condition variability for the `ImageNet` images. We observed that this was because agreement was driven by a subset of adversarial images (see Supplementary Figure 2.3 for a break down). Thus, even for these `ImageNet` images, DCNN representations do not consistently overlap with representations used by humans.

### 2.6. Experiment 4: Generating fooling images for ImageNet

Experiment 3 showed that it is straightforward to obtain overall chance level performance on the `MNIST` images, and this raises the obvious question of whether it is also straightforward to observe chance performance for adversarial images designed to fool `ImageNet` classifiers? In order to test this we generated our own irregular (TV-static like) adversarial images using a standard method of generating adversarial images (see Methods section). Each of these images was confidently classified as one out of a 1000 categories by a network trained on `ImageNet`. Participants were presented three of these adversarial images and asked to choose the image that most closely matches the target category (see inset in Figure 7). In half of the trials participants were shown adversarial images that were generated to fool `AlexNet` while in the other half they were shown adversarial images generated to fool `Resnet-18`.

**Figure 7:**
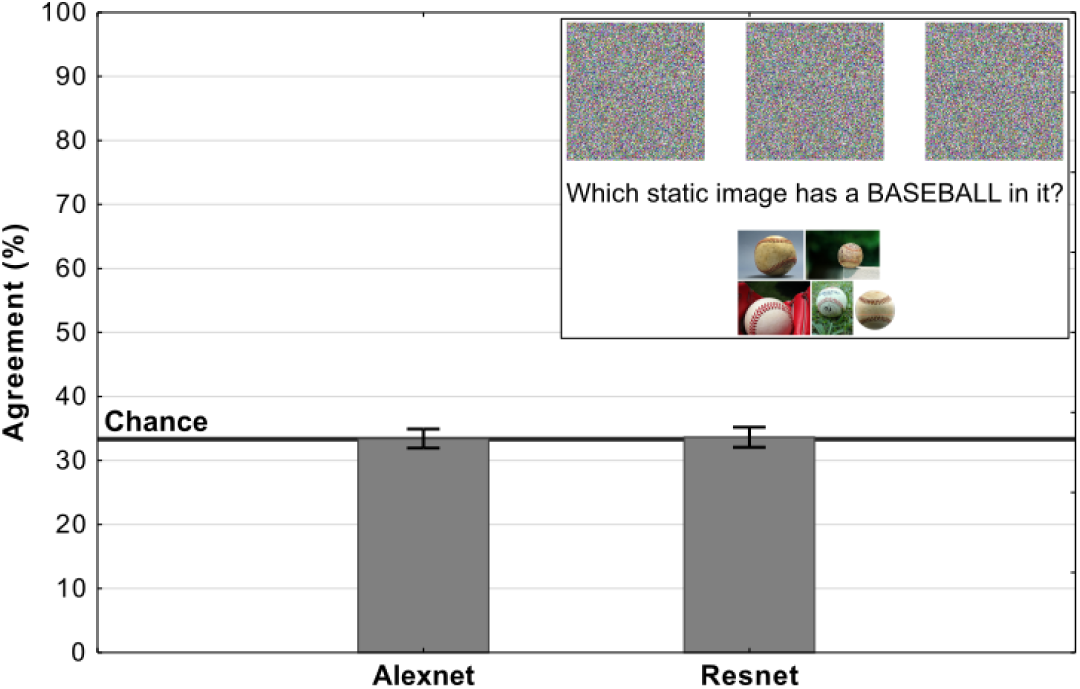
Average levels of agreement in Experiment 4 (error bars denote 95% confidence intervals). The inset depicts a single trial in which participants were shown three fooling adversarial images and naturalistic examples from the target category. Their task was to choose the adversarial image which contained an object from the target category.

Results of the experiment are shown in Figure 7. For both types of images, the agreement between participants and DCNNs was at chance. Additionally, we ran binomial tests for each image in order to determine whether the number of participants which agreed with DCNN classification was significantly above chance and the results showed that not a single image showed agreement that was significantly above chance. Clearly, participants could not find meaningful features in any of these images, while networks were able to confidently classify each of these images.

## 3. Discussion

Adversarial images may look meaningless at first glance, but they contain features that convolutional networks find highly diagnostic of an image category. If human classification of these images strongly correlates with DCNNs, as Zhou and Firestone [37] observed, it would suggest that there may be common mechanistic processes underlying human and DCNN object recognition.

However, when we examined their results more carefully, we found that the level of agreement was much lower than reported and largely driven by a subset of images where participants could eliminate response alternatives based on some superficial features present within these images. This was confirmed in a series of experiments that found that agreement between humans and DC-NNs was contingent on the adversarial images chosen as stimuli (Experiments 2 and 3) and the response alternative presented to participants (Experiment 1). This suggests that humans and current architectures of DCNNs process adversarial images in fundamentally different ways: while humans are making educated guesses based on some superficial features (colour, texture, curvature, etc.) of these images, current architectures of DCNNs are relying on highly diagnostic features that humans are ignoring in order to support high-confidence judgements. Furthermore, we also show how easy it is to generate adversarial images that lead to overall chance level DCNN-human agreement (Experiments 3 and 4), again highlighting the qualitative differences in these systems, with DCNNs confidently identifying images on the basis of features that humans completely ignore.

A similar distinction between human and DCNN classification is made by Ilyas et al. [20], who argue that current architectures of DCNNs are vulnerable to adversarial attacks due to their tendency for relying on *non-robust features* present in databases. These are features that are predictive of a category but highly sensitive to small perturbations of the image. It is this propensity for relying on non-robust features that makes it easy to generate adversarial images that are completely uninterpretable by humans but classified confidently by the network (Experiment 4). A striking example of DCNNs picking up on non-robust features was recently reported by Malhotra and Bowers [25] who showed that DCNNs trained on a `CIFAR-10` dataset modified to contain a single diagnostic pixel per category, learn to categorize images based on single pixels ignoring everything else in the image. Humans, by contrast, tend to use robust features of objects, such as their shape, for classifying images [8].

We would like to note that we are not claiming that there is no role played by superficial and non-robust features in human object recognition. In a recent study, [15] asked human participants to classify naturalistic adversarial images (see Figure 1(b)) when these images were briefly flashed (for around 70ms) on the screen. They found that there is a small, but statistically significant, effect of the adversarial manipulation on choices made by participants (i.e., participants were slightly more likely to classify a ‘cat’ image as a ‘dog’ when the image was adversarially perturbed towards a ‘dog’). Thus, these results seem to suggest that humans are sensitive to the same type of non-robust features that lead to adversarial attacks on DCNNs. However, it is important to note here that the size of these effects is small: while human accuracy drops by less than 10% when normal images are replaced by adversarially perturbed images, DCNNs (mis)classify these adversarially perturbed images with high confidence. These findings are consistent with our observation that some adversarial images capture some superficial features that can be used by participants to make classification decisions, leading to an overall above-chance agreement.

It should also be noted that we have only considered a small fraction of adversarial images here and, like Experiment 4, there are many other types of adversarial attacks that produce images that seem completely undecipherable for humans. It could be that humans find these images completely uninterpretable due to the difference in *acuity* of human and machine vision (a line taken by *Z&F*). There are two reasons why we think a difference in acuity cannot be the primary explanation of the difference between human and machine perception of adversarial images. Firstly, we have shown above that the very same algorithm produced some images that supported above chance agreement and other images that supported no agreement (for example, Supplementary Figure 2.2). There is no reason to believe that the two sets of images are qualitatively different, with DCNNs selectively exploiting subliminal features only when overall agreement levels are chance. Secondly, a wide variety of adversarial attacks clearly do not rely on subtle visual features that are below human perceptual threshold. This includes semantic adversarial attacks that occur when the colour of an images is changed [19], or attacks that cause incorrect classification by simply change the pose of an image [2], etc. These are all dramatic examples of differences between DCNNs and humans that cannot be attributed to the acuity of human perceptual front-end. Rather they reflect the fact that current architectures of DCNNs are often relying on visual features that humans can see but ignore.

To conclude, our findings with fooling adversarial images highlight how DC-NNs are currently poor psychological models of human object identification. A key challenge for future research is to develop models that are sensitive to the visual features that humans rely on but at the same time insensitive to other features that are diagnostic of object category but irrelevant to human vision. This involves identifying objects on the basis of shape rather than texture or color [16, 4], where vertices are the critical components of images [5], where Gestalt principles are used to organize features [30], where relations between parts are explicitly coded [34], where features and objects are coded independently of retinal position [9] size [7], left/right reflection [6], etc. When DCNNs rely on these set of features, we expect they will not be subject to adversarial attacks that seem so bizarre to humans, and will show the same set of of strengths and weakness (visual illusions) that characterize human vision.

## 4. Methods

### Reassessing agreement

**Blindfolded participants** If a participant is blind-folded and chooses one of 48 options randomly on 48 trials, the probability of them making the same choice as the DCNN on *k* trials is given by the binomial distribution 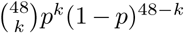, where 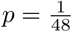. Substituting different values of *k*, one can compute that 37.2% of these blindfolded participants will agree with the DCNN on 1 trial, 18.6% will agree on 2 trials, 6% will agree on 3 trials, and so on. To compute the proportion of participants who agree with the DCNN, Zhou and Firestone [37] count all participants who agree on 2 or more trials as agreeing with the DCNN (chance is 1 out of 48 trials) and half of the participants that agree on exactly 1 trial. Thus, summing up all the blindfolded participants that agree on 2 or more trials and half of those who agree on exactly 1 trial, this method will show ∼ 45% agreement between participants and the DCNN. **Experiment 1.** This experiment examined whether agreement between humans and DCNNs depended on the response alternatives presented to participants. We tested *N* = 200 participants and each participant completed 10 trials. During each trial, participants were presented a fooling adversarial image and four response alternatives underneath the image and asked to choose one of these alternatives. Participants indicated their response by moving their cursor to the response alternative and clicking. We selected 10 fooling adversarial images from amongst the 48 images used by *Z&F* in in their Experiments 1–3. Each of these images was classified with *>* 99% confidence by a DCNN which was trained to classify the `ImageNet` dataset. We selected these 10 adversarial images to minimise semantic and functional overlap in the labelled categories (for example, we avoided selecting both ‘computer keyboard’ and ‘remote control’). The experiment consisted of two conditions, which differed in how the response alternatives were chosen on each trial. In the ‘Competitive’ condition (*N* = 100) we chose four response alternatives that subjectively seemed to contain one or more visual features that were present in the adversarial image. One of these response alternatives was always the label chosen by the DCNN. The other three were chosen from amongst 1000 `ImageNet` class labels. This was again done to minimise a semantic or functional overlap with the target class (e.g. an alternative for the ‘baseball’ class was *parallel bars* but not *basketball*). All ten images and the four competitive response alternatives for each image are shown in Supplementary Figure 2.1. In the ‘Random’ condition (*N* = 100) the three remaining alternative responses were drawn at random (on each trial) from the aforementioned 48 target classes from Experiment 3 in Zhou and Firestone [37]. We randomised the order of images, as well as the order of the response alternatives for each participant.

### Experiment 2

This experiment was designed to examine whether all fooling adversarial images for a category show similar levels of agreement between humans and DCNNs. The experiment’s design was the same as Experiment 1 above, except participants were now randomly assigned to the ‘best-case’ (*N* = 101) and ‘worst-case’ (*N* = 100) conditions. In each condition, participants again completed 10 trials and on each trial, they saw an adversarial image and four response alternatives. One of these alternatives was the category chosen by a DCNN with > 99% confidence and the other three were randomly drawn from amongst the 48 categories used by *Z&F* in their Experiments 1–3. The difference between the ‘best-case’ and ‘worst-case’ conditions was the adversarial image that was shown to the participants on each trial. Each of the adversarial images used by *Z&F* in their Experiments 1–3 were chosen from the set of images generated by Nguyen et al. [26]. In that paper, Nguyen et al. [26] generated 5 adversarial images for each `ImageNet` category. In the ‘best-case’ condition, we selected an adversarial image for each category (from amongst these 5 alternatives) that subjectively seemed to contain the largest number of superficial visual features present in examples of the target category. Similarly, in the ‘worst-case’ condition, we selected an adversarial image for each category that seemed to contain the least number of superficial visual features in common with objects of the target category. DCNNs showed the same confidence in classifying both sets of images. We again randomised the order of presentation of images.

### Experiment 3

The experiment consisted of four experimental conditions in a 2×2 repeated measures design (every participant completed each condition). The first factor of variation was the database on which the DCNNs were trained − `ImageNet` or `MNIST` with one condition containing images designed to fool `ImageNet` and the other containing images designed to fool `MNIST` classifiers. The second factor of variation was the evolutionary algorithm used to generate the adversarial images – *direct* or *indirect*. The indirect encoding method leads to adversarial images which have regular features (e.g. edges) that often repeat, while the direct encoding method leads to noise-like adversarial images. All of the images were from the seminal Nguyen et al. [26] paper on fooling images. The `MNIST` dataset consists of ten categories (corresponding to handwritten numbers between 0 and 9), while `ImageNet` consists of 1000 categories. As we wanted to compare agreement across conditions, we selected ten images (from ten different categories) for both datasets. The indirectly-encoded `ImageNet` images were the same as the ones in Experiment 1 while the images for the other three conditions were randomly sampled from the images generated by [26]. Participants were shown one image at a time and asked to categorize it as one of ten categories (category labels were shown beneath the image). One of these ten categories was the label assigned to the image by a DCNN. Therefore, chance level agreement was 10%. The participants had to click on the label they thought represented what was in the image. The order of conditions was randomized for each participant and the order of images within each condition was randomized as well. A total of *N* = 200 participants completed the study. Two participants were excluded from analysis because their choices were made with average response times below 500 ms indicating random clicking rather than actually making decisions based on looking at the images themselves.

### Experiment 4

In this experiment we used the Foolbox package ([32]) to generate images that fool DCNNs trained on `ImageNet`. The experiment consisted of two conditions, one with images designed to fool AlexNet [23] and the other with images designed to fool ResNet-18, both trained on `ImageNet`. We generated our own adversarial images by first generating an image in which each pixel was independently sampled and successively modifying this image using an Iterative Gradient Attack based on the fast gradient sign method ([17]) until a DCNN classified this image as a target category with a > 99% confidence. The single trial procedure mirrors Experiment 4 from *Z&F*. Participants (*N* = 200) were shown three of the generated images and a set of five real-world example images of the target class (see Inset in Figure 7). They were asked to choose the adversarial image which contained an object from the target class. The example images were randomly chosen from the `ImageNet` dataset for each class. Each participant completed both experimental conditions. The order of trials was randomized for each participant.

### Statistical analyses

All conducted statistical analyses were two-tailed with a p-value under 0.05 denoting a significant result. For the reanalysis of data from [37] we used Binomial tests. In Experiments 1, 2, and 4 we conducted single sample t-tests to check if agreement levels were significantly above a fixed chance level (25% in Experiments 1 and 2, 33.33% in Experiment 4). We additionally ran a between-subject t-test (Experiments 1 and 2) and a within-subject t-test (Experiment 4) to determine whether the difference between experimental conditions was significant. We also conduct a Binomial test in Experiment 4 to determine for how many items was agreement level significantly above chance. In Experiment 3 we ran a two-way repeated measures analysis of variance. We report effect size measures for all tests (Cohen’s d for t-tests and partial eta squared for ANOVA effects).

### Power analysis

A sample size of *N* = 200 was chosen for each experiment which mirrors *Z&F* experiments 1-6 in order to detect similar effects. This allowed us to detect an effect size as low as *d* = 0.18 at *α* = .05 with 0.80 power in within-subject and *d* = 0.35 in between-subject experiments.

### Online recruitment

We conducted all four experiments online with recruitment through the Prolific platform. Each sample was recruited from a pool of registered participants which met the following criteria. Fluent English speakers living in the UK, USA, Canada or Australia of both genders between the ages of 18 and 50 with normal or corrected to normal vision and a high feedback rating on the Prolific platform (above 90). Participants were reimbursed for their time upon successful completion through the Prolific system.

### Data availability

Data and stimuli are available via the Open Science Framework at https://osf.io/a2sh5/.

### Ethics approval

This study was approved by the Research Ethics Committee of the University of Bristol.

## 5. Acknowledgments

This research was supported by the European Research Council Grant Generalization in Mind and Machine, ID number 741134. We would like to thank Alex Doumas, Jeff Mitchell, Milton Liera Montero and Brian Sullivan for their insights and feedback.

