## Supplementary materials for "What do adversarial images tell us about human vision?"

### 1. Reanalysis of Zhou and Firestone (2019)

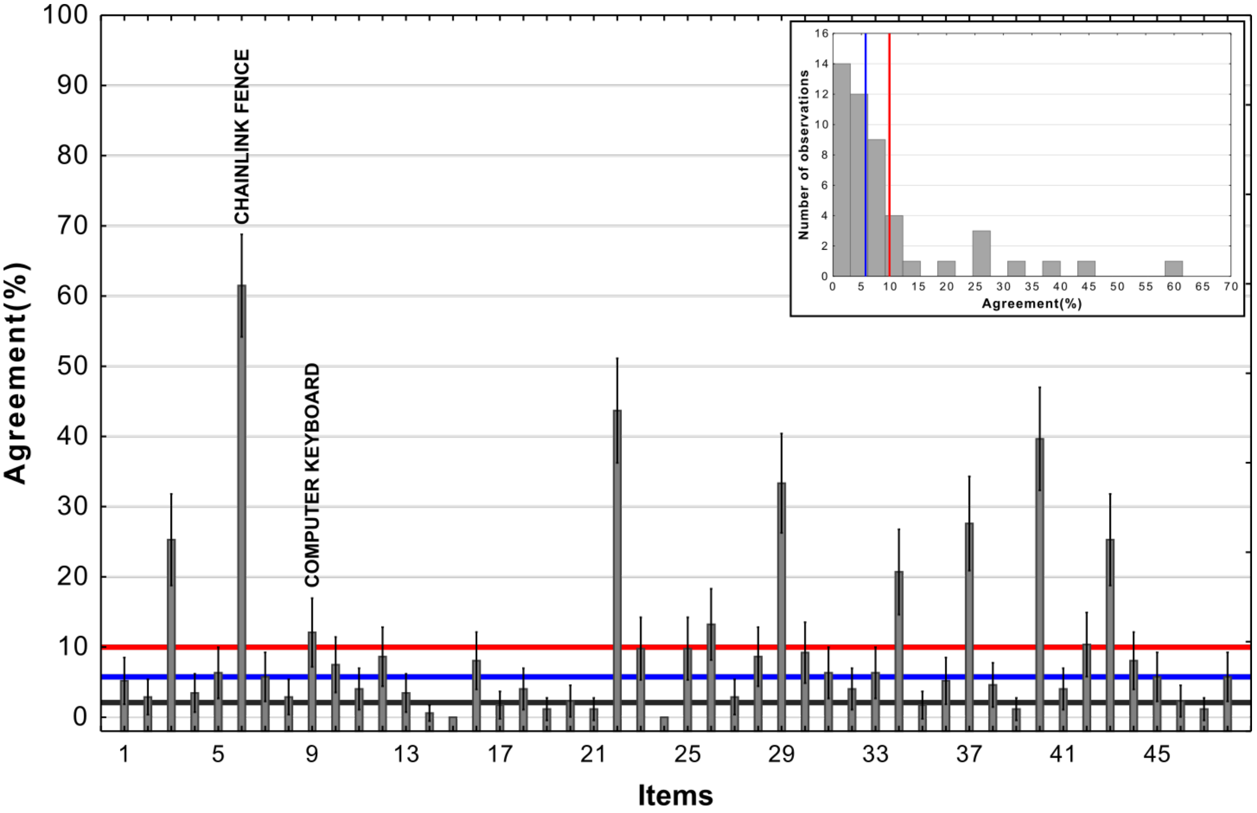


**Supplementary Figure 1.1: Agreement across adversarial images from Experiment 3b in Zhou and Firestone (2019)** The red line represents the mean, the blue line represents the median, and the black reference line represents chance agreement. The inset contains a histogram of agreement levels across the 48 images.

**Supplementary discussion 1.1**

Roughly similar agreement levels on most images accompanied by mean agreement above chance would indicate some systematic underlying overlap between human and network object recognition. However, as shown in Supplementary Figure 1.1, there are vast differences in agreement levels depending on adversarial image. The inset of Supplementary Figure 1.1 also shows that the distribution is skewed and that the mean agreement metric overestimates true human-network agreement which is better represented by the median. There is a minority of images which drive agreement as outliers, giving credence to the hypothesis about two separate sources of agreement discussed in the paper.

| Stimulus | DCNN label | 1 | 2 | 3 | 4 | 5 | 6 | 7 | 8 | | |
|---|---|---|---|---|---|---|---|---|---|---|---|
| 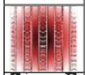 | 1. Accordion** | Traffic light 10.92% | Strawberry 8.05% | Tile roof 6.32% | **Accordion 5.17%** | Spotlight 5.17% | Electric Guitar 4.60% | Pole 4.60% | Chainlink fence 4.02% | | |
| 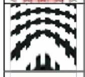 | 2. Assault rifle* | Chainlink fence 21.26% | Crossword puzzle 8.04% | Computer keyboard 6.32% | Accordion 5.17% | Soccer ball 4.60% | Ski mask 3.45% | Tile roof 3.45% | **Assault rifle 2.87%** | | |
| 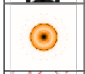 | 3. Bagel** | **Bagel 25.28%** | Spotlight 12.07% | Traffic light 5.74% | Volcano 5.17% | Pinwheel 4.02% | Car wheel 2.87% | Chameleon 2.87% | Digital clock 2.87% | | |
| 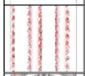 | 4. Baseball* | Pole 12.64% | Four-poster bed 5.74% | Sea snake 5.74% | Chainlink fence 4.59% | Hair Clip 4.02% | Panpipe 4.02% | Roundworm 4.02% | **Baseball 3.45%** | | |
| 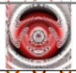 | 5. Car Wheel** | Traffic light 10.92% | Volcano 6.90% | **Car wheel 6.32%** | Obelisk 5.75% | Stethoscope 5.75% | Electric guitar 4.02% | Projector 4.02% | Digital clock 3.45% | | |
| 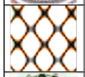 | 6. Chainlink fence** | **Chainlink fence 61.49%** | Accordion 2.87% | Hair clip 2.30% | Sea snake 2.30% | Ski mask 2.30% | Tile roof 2.30% | Computer keyboard 1.72% | Digital clock 1.72% | | |
| 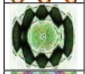 | 7. Chameleon** | Pinwheel 6.90% | **Chameleon 5.75%** | Starfish 5.75% | Green snake 5.17% | Stethoscope 5.17% | Peacock 4.60% | Projector 4.60% | Traffic light 4.60% | | |
| 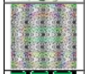 | 8. Comic book* | Peacock 8.05% | Projector 6.90% | Chainlink fence 5.75% | Chameleon 5.75% | Green snake 4.60% | Remote control 4.60% | Accordion 4.02% | Stethoscope 4.02% | … | **Comic book (11) 2.87%** |
| 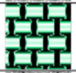 | 9. Computer keyboard** | Chainlink fence 14.37% | **Computer keyboard 12.07%** | Green snake 8.05% | Tile roof 4.60% | Paddle 4.02% | Chameleon 3.45% | Digital clock 3.45% | Slot machine 3.45% | | |
| 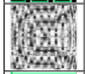 | 10. Crossword puzzle** | Chainlink fence 8.62% | **Crossword puzzle 7.45%** | Tile roof 6.90% | Computer keyboard 6.32% | Accordion 4.60% | Electric guitar 4.60% | Photocopier 4.60% | Digital clock 4.02% | | |
| 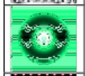 | 11. Dial telephone* | Traffic light 9.77% | Soccer ball 8.05% | Bagel 6.32% | Car wheel 6.32% | Chameleon 6.32% | Projector 5.75% | Baseball 5.17% | Roundworm 4.60% | … | **Dial telephone (10) 4.02%** |
| 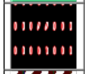 | 12. Digital clock** | Strawberry 14.37% | Roundworm 12.07% | **Digital clock 8.62%** | Accordion 4.60% | Remote control 4.60% | Slot machine 4.60% | Spotlight 4.02% | Computer keyboard 3.45% | | |
| 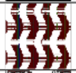 | 13. Electric guitar* | Sea snake 12.07% | Chainlink fence 6.32% | Hair clip 5.75% | Roundworm 5.75% | Tile roof 5.75% | Accordion 4.02% | Hand blower 4.02% | **Electric guitar 3.45%** | | |
| 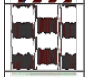 | 14. Four-poster bed | Accordion 8.62% | Tile roof 7.47% | Freight car 5.17% | Chainlink fence 4.60% | Soccer ball 4.60% | Electric guitar 4.02% | Paddle 4.02% | Comic book 3.45% | … | **Four poster bed (35) 0.57%** |
| 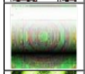 | 15. Freight car | Projector 8.05% | Peacock 7.47% | Digital clock 6.90% | Electric guitar 6.90% | Stethoscope 6.90% | Photocopier 5.75% | Slot machine 5.17% | Chameleon 4.60% | … | **Freight car (39) 0%** |
| 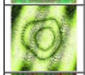 | 16. Green snake** | **Green snake 8.05%** | Roundworm 8.05% | Spotlight 8.05% | Chameleon 6.90% | Bagel 4.02% | Digital clock 4.02% | Soccer ball 4.02% | Traffic light 4.02% | | |
| 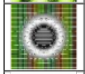 | 17. Grey parrot | Bagel 14.37% | Stethoscope 8.05% | Car wheel 5.75% | Soccer ball 5.17% | Spotlight 5.17% | Vacuum 5.17% | Baseball 4.60% | Projector 4.60% | … | **Grey parrot (15) 1.72%** |
| 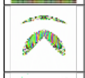 | 18. Hair clip* | Monarch butterfly 32.76% | Peacock 6.32% | Chameleon 5.75% | Ski mask 5.75% | Obelisk 4.60% | **Hair clip 4.02%** | Green snake 3.45% | Paddle 2.87% | | |
| 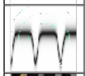 | 19. Hand blower | Computer keyboard 9.20% | Digital clock 6.90% | Stethoscope 5.17% | Volcano 4.60% | Accordion 4.02% | Hair clip 4.02% | Chameleon 3.45% | Dial telephone 3.45% | … | **Hand blower (25) 1.15%** |
| 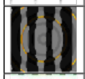 | 20. King penguin* | Four-poster bed 7.47% | Car wheel 5.75% | Chainlink fence 4.60% | Stethoscope 4.60% | Projector 4.02% | Grey parrot 3.45% | Punching bag 3.45% | Slot machine 3.45% | … | **King penguin (12) 2.30%** |
| 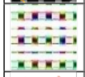 | 21. Medicine chest | Slot machine 8.62% | Computer keyboard 8.05% | Chainlink fence 6.32% | Photocopier 6.32% | Electric guitar 5.17% | Panpipe 5.17% | Chameleon 4.60% | Tile roof 4.60% | … | **Medicine chest (20) 1.15%** |
| 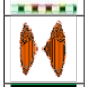 | 22. Monarch butterfly** | **Monarch butterfly 43.68%** | Volcano 8.05% | Punching bag 4.02% | Hair clip 3.45% | Paddle 3.45% | Starfish 3.45% | Stethoscope 3.45% | Traffic light 3.45% | | |
| 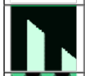 | 23. Obelisk** | **Obelisk 9.77%** | Pole 6.90% | Four-poster bed 4.60% | Projector 4.60% | Tile roof 4.60% | Chainlink fence 4.02% | Crossword puzzle 4.02% | Photocopier 4.02% | | |
| 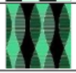 | 24. Paddle | Green snake 25.86% | Sea snake 6.32% | Accordion 4.02% | Obelisk 4.02% | Chainlink fence 3.45% | Chameleon 3.45% | Electric guitar 3.45% | Hand blower 3.45% | … | **Paddle (41) 0%** |

* numerically above chance agreement; ** statistically above chance agreement

**Supplementary Figure 1.2: Participant responses ranked by frequency (Experiment 3b).** Each row contains the adversarial image, the DCNN label for that image, the top 8 participant responses. Shaded cells contain the DCNN choice, when not ranked in the top 8, it is shown at the end of the row along with the rank in brackets.

| Timulus | DCNN label | 1 | 2 | 3 | 4 | 5 | 6 | 7 | 8 | | |
|---|---|---|---|---|---|---|---|---|---|---|---|
| 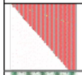 | 25. Panpipe** | Tile roof 12.07% | **Panpipe 9.77%** | Accordion 4.60% | Obelisk 4.02% | Hair clip 3.45% | Hand blower 3.45% | Pinwheel 3.45% | Pole 3.45% | | |
| 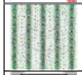 | 26. Peacock** | Green snake 13.22% | **Peacock 13.22%** | Sea snake 7.47% | Accordion 5.75% | Chainlink fence 5.75% | Monarch butterfly 4.02% | Chameleon 3.45% | Crossword puzzle 3.45% | | |
| 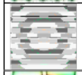 | 27. Photocopier* | Car wheel 16.67% | Spotlight 6.32% | Pinwheel 4.60% | Accordion 4.02% | Projector 4.02% | Traffic light 4.02% | Assault rifle 3.45% | Digital clock 3.45% | ... | **Photocopier (11) 2.87%** |
| 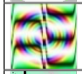 | 28. Pinwheel** | **Pinwheel 8.62%** | Computer keyboard 6.90% | Projector 6.32% | Chameleon 5.75% | Peacock 5.17% | Slot machine 4.60% | Spotlight 4.60% | Monarch butterfly 4.02% | | |
| 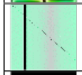 | 29. Pole** | **Pole 33.33%** | Traffic light 5.17% | Comic book 4.02% | Dial telephone 4.02% | Obelisk 4.02% | Photocopier 4.02% | Chainlink fence 3.45% | Accordion 2.87% | | |
| 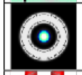 | 30. Projector** | Spotlight 19.54% | Car wheel 9.20% | **Projector 9.20%** | Stethoscope 5.75% | Soccer ball 5.17% | Traffic light 4.02% | Assault rifle 3.45% | Digital clock 3.45% | | |
| 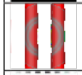 | 31. Punching bag** | Pole 20.69% | **Punching bag 6.32%** | Spotlight 5.75% | Remote control 4.02% | Roundworm 4.02% | Traffic light 4.02% | Four-poster bed 3.45% | Accordion 2.87% | | |
| 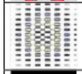 | 32. Remote control* | Computer keyboard 10.92% | Chainlink fence 8.05% | Tile roof 5.75% | Stethoscope 5.17% | Digital clock 4.60% | Projector 4.60% | Obelisk 4.02% | **Remote control 4.02%** | | |
| 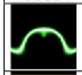 | 33. Roundworm** | Green snake 27.01% | **Roundworm 6.32%** | Sea snake 5.75% | Hair clip 5.75% | Electric guitar 3.45% | Digital clock 3.45% | Ski mask 3.45% | Slot machine 3.45% | | |
| 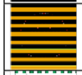 | 34. School bus** | **School bus 20.69%** | Accordion 5.17% | Photocopier 5.17% | Chainlink fence 4.02% | Monarch butterfly 4.02% | Medicine chest 3.45% | Tile roof 3.45% | Chameleon 2.87% | | |
| 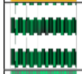 | 35. Screwdriver | Accordion 12.64% | Panpipe 9.77% | Green snake 8.62% | Chainlink fence 5.75% | Computer keyboard 5.75% | Digital clock 4.02% | Slot machine 4.02% | Stethoscope 3.45% | ... | **Screwdriver (17) 1.72%** |
| 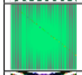 | 36. Sea snake** | Traffic light 14.37% | Green snake 9.20% | Pole 6.90% | Spotlight 5.75% | Chameleon 5.17% | **Sea snake 5.17%** | Digital clock 4.60% | Photocopier 4.60% | | |
| 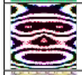 | 37. Ski mask** | **Ski mask 27.59%** | King penguin 8.05% | Monarch butterfly 8.05% | Peacock 4.60% | Electric guitar 3.45% | Pinwheel 3.45% | Chameleon 2.87% | Comic book 2.87% | | |
| 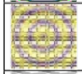 | 38. Slot machine* | Spotlight 10.35% | Pinwheel 9.20% | Crossword puzzle 5.17% | Projector 5.17% | Bagel 4.60% | **Slot machine 4.60%** | Traffic light 4.60% | Computer keyboard 4.02% | | |
| 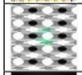 | 39. Soccer ball | Chainlink fence 17.24% | Crossword puzzle 6.90% | Photocopier 6.32% | Electric guitar 5.75% | Accordion 5.17% | Traffic light 4.60% | Digital clock 2.87% | Projector 2.87% | ... | **Soccer ball (26) 1.15%** |
| 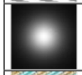 | 40. Spotlight** | **Spotlight 39.66%** | Projector 15.52% | Traffic light 5.75% | Vacuum 4.60% | Obelisk 3.45% | Stethoscope 3.45% | Soccer ball 2.87% | Freight car 2.30% | | |
| 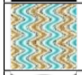 | 41. Starfish* | Sea snake 11.49% | Electric guitar 6.90% | Peacock 6.90% | Accordion 6.32% | Chainlink fence 5.17% | Slot machine 4.60% | Pinwheel 4.02% | **Starfish 4.02%** | | |
| 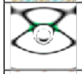 | 42. Stethoscope** | Car wheel 12.06% | **Stethoscope 10.34%** | Chainlink fence 6.90% | Spotlight 6.32% | Obelisk 4.02% | Photocopier 4.02% | Projector 4.02% | Soccer ball 4.02% | | |
| 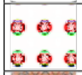 | 43. Strawberry** | **Strawberry 25.29%** | Traffic light 16.67% | Slot machine 6.90% | Baseball 4.60% | Monarch butterfly 3.45% | Stethoscope 3.45% | Soccer ball 2.87% | Medicine chest 2.30% | | |
| 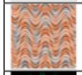 | 44. Tile roof** | Accordion 8.62% | Volcano 8.62% | **Tile roof 8.05%** | Monarch butterfly 6.90% | Chainlink fence 5.17% | Starfish 5.17% | Dial telephone 4.02% | Hand blower 4.02% | | |
| 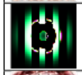 | 45. Traffic light** | Projector 7.47% | Pinwheel 6.90% | Spotlight 6.32% | Car wheel 5.75% | **Traffic light 5.75%** | Stethoscope 4.02% | Obelisk 3.45% | Photocopier 3.45% | | |
| 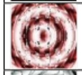 | 46. Trifle* | Strawberry 9.77% | Pinwheel 7.47% | Roundworm 6.90% | Traffic light 6.90% | Spotlight 6.32% | Volcano 5.75% | Car wheel 4.60% | Stethoscope 4.60% | ... | **Trifle (13) 2.30%** |
| 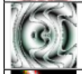 | 47. Vacuum | Green snake 4.60% | Chameleon 4.02% | Monarch butterfly 4.02% | Obelisk 4.02% | Peacock 4.02% | Pinwheel 4.02% | Projector 4.02% | Starfish 4.02% | ... | **Vacuum (31) 1.15%** |
| 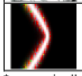 | 48. Volcano** | Pole 9.77% | Hair clip 6.90% | Spotlight 6.90% | Traffic light 6.32% | **Volcano 5.75%** | Roundworm 5.17% | Panpipe 4.60% | Punching bag 4.60% | | |

* numerically above chance agreement; ** statistically above chance agreement

**Supplementary Figure 1.2: Participant responses ranked by frequency (Experiment 3b).** Continued.

**Supplementary discussion 1.2**

When trying to determine the nature of human-network agreement it is important to consider distributions of choices on a per-item level not merely whether agreement was above chance for a particular stimulus. If there was general agreement between humans and networks one could expect that for most images the most frequently chosen label would be the one assigned to the stimulus by the network. Additionally, it could be expected that a large percentage of participants choose exactly this label. Supplementary Figure 1.2 shows the top eight most frequently chosen labels by participants in Experiment 3b from Zhou and Firestone (2019). It can clearly be seeen that for only a minority of images the network label was the most often chosen one by participants. Additionally, there is only one stimulus for which majority of participants chose the network label (e.g. 'chainlink fence', image number 6 in the Supplementary Table 1) while only for a few others can it be said that a fairly substantial percentage of participants chose the network label. For many of the stimuli, the network label is not amongst the top eight choices made by participants. It can also be observed that for most stimuli the top most frequently made choice wasn't overwhelmingly favoured by participants, meaning the distribution of label choices for most stimuli is flat, indicating guessing rather than agreement (see Supplementary Figure 1.3 for more information).

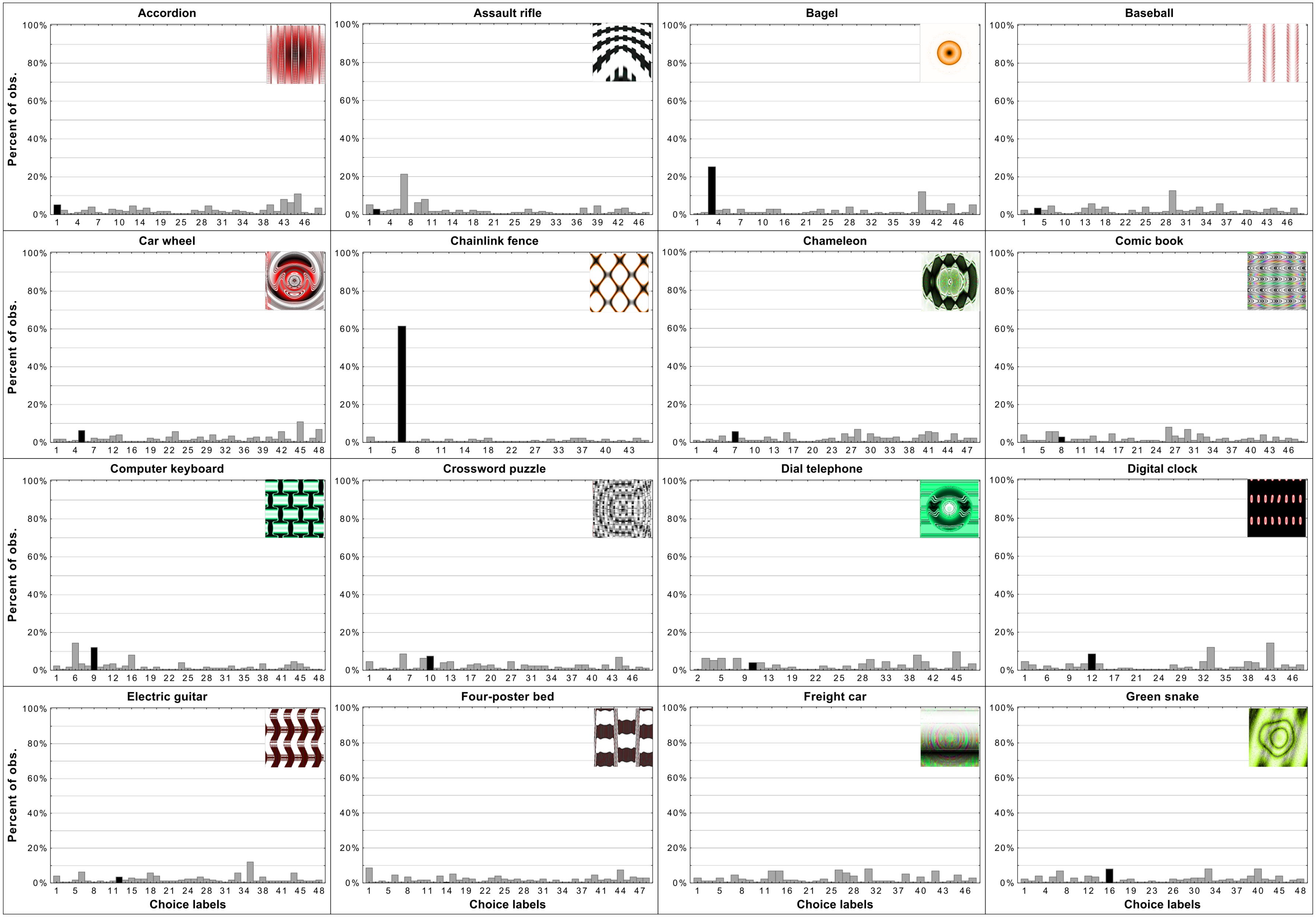


**Supplementary Figure 1.3: Per-item histograms of response choices from Experiment 3b in Zhou and Firestone(2019)** Each histogram contains the adversarial stimuli and shows the percentage of responses per each choice (y-axis). The choice labels (x-axis) are ordered the same way as in Supplementary Table 1 from 1 to 48. Black bars indicate the DCNN choice for a particular adversarial image.

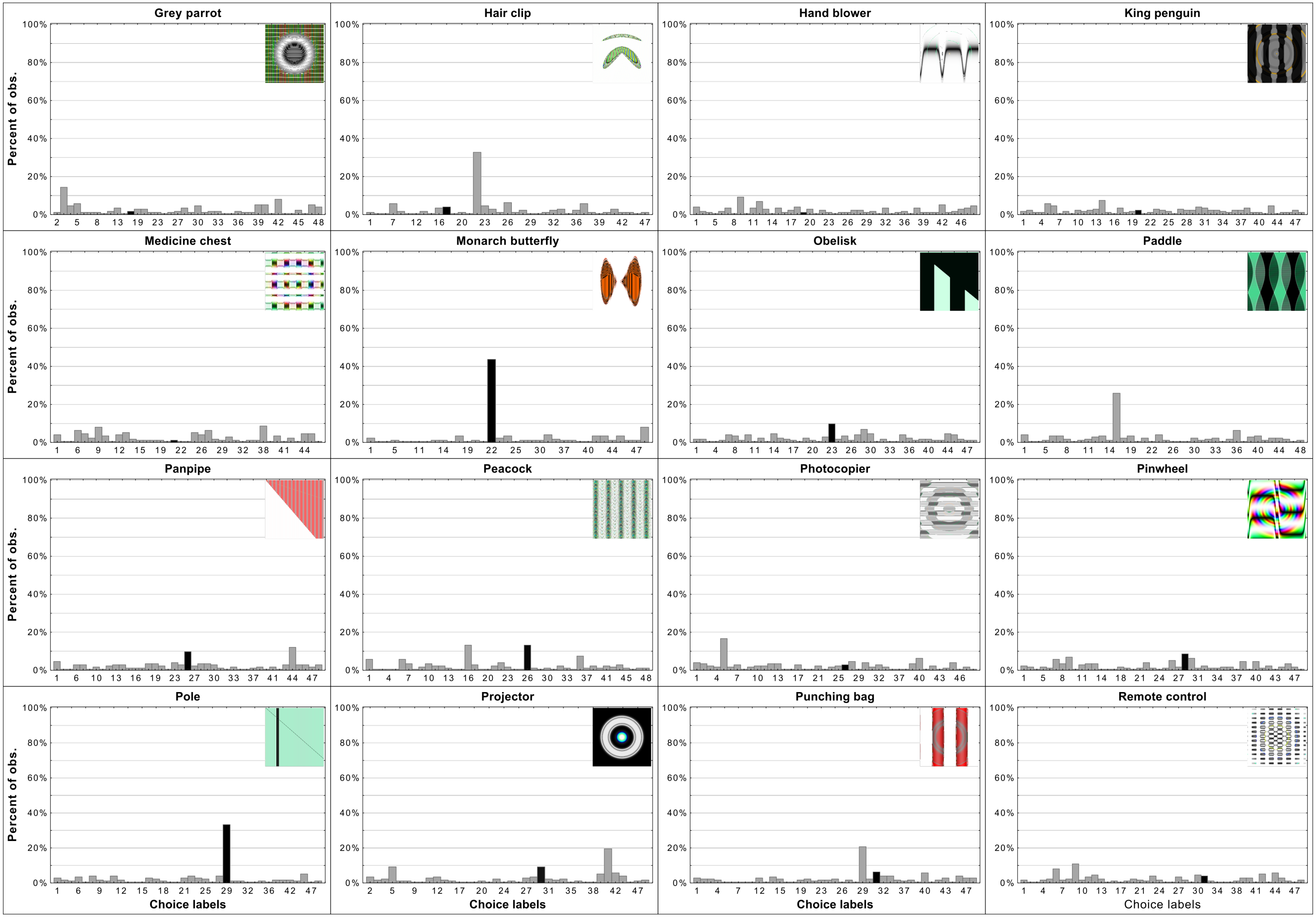


**Supplementary Figure 1.3: Per-item histograms of response choices from Experiment 3b in ?.** Continued.

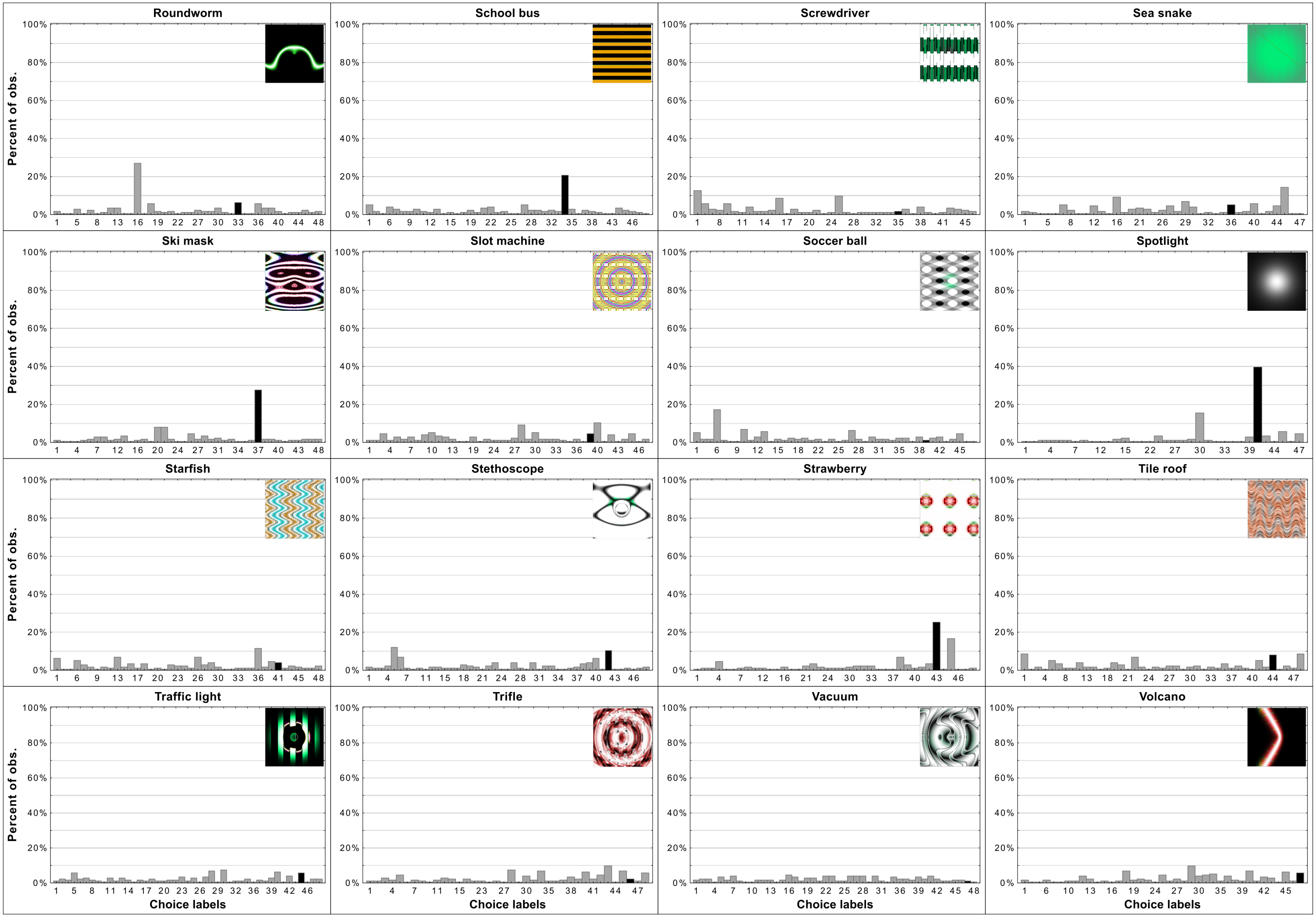


**Supplementary Figure 1.3: Per-item histograms of response choices from Experiment 3b in ?.** Continued.

**Supplementary discussion 1.3**

As was implied in Supplementary Figure 1.2, Supplementary Figure 1.3 reveals flat distributions of label choices for the vast majority of stimuli. Indeed only eight histograms resemble what could be expected if there was systematic human-network agreement on classification. For those stimuli the most often chosen label was the network label and the percentage of participants who chose the label peaks above the percentage of choices for other labels. There are examples with similar peaks but in which the most often chosen label was not the one assigned to the stimulus by DCNNs. Overall, this provides evidence for the hypothesis that agreement is derived from two sources. First, some stimuli (e.g. 'chainlink fence') can hardly be called adversarial images since they retain almost all of the features as well as maintain the relationship between features of the target category. In those rare cases, agreement is trivial. In other cases in which agreement is above chance it is likely that those levels were achieved by excluding labels based on a few superficial features. These features were not sufficient for object recognition but did allow for exclusion of labels which do not contain the specific features (e.g. dismissing 'monarch butterfly' when viewing the 'projector' stimulus). We believe that this exact pattern of data supports such a hypothesis.

### 2. Supplementary information for Experiments 1-4

| Stimulus | DCNN label | Alternative 1 | Alternative 2 | Alternative 3 |
| --- | --- | --- | --- | --- |
| 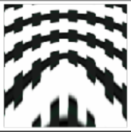 | Assault rifle | Accordion | Chainsaw | Piano |
| 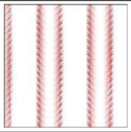 | Baseball | Bath towel | Parallel bars | Wool |
| 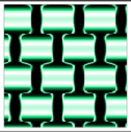 | Computer keyboard | Chainlink fence | Crossword puzzle | Honeycomb |
| 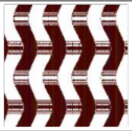 | Electric guitar | Harmonica | Picket fence | Radiator grille |
| 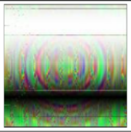 | Freight car | Bubble | Car wheel | Traffic light |
| 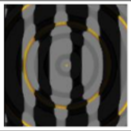 | King penguin | CD player | Plate | Wall clock |
| 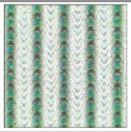 | Peacock | Broccoli | Grass snake | Valley |
| 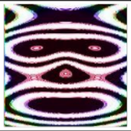 | Ski mask | Baboon | Loudspeaker | Totem pole |
| 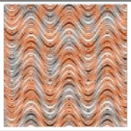 | Tile roof | Rock crab | Shower curtain | Wooden spoon |
| 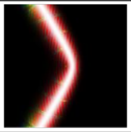 | Volcano | Lighter | Missile | Table lamp |

**Supplementary Figure 2.1: Experiment 1 stimuli and alternative labels.**

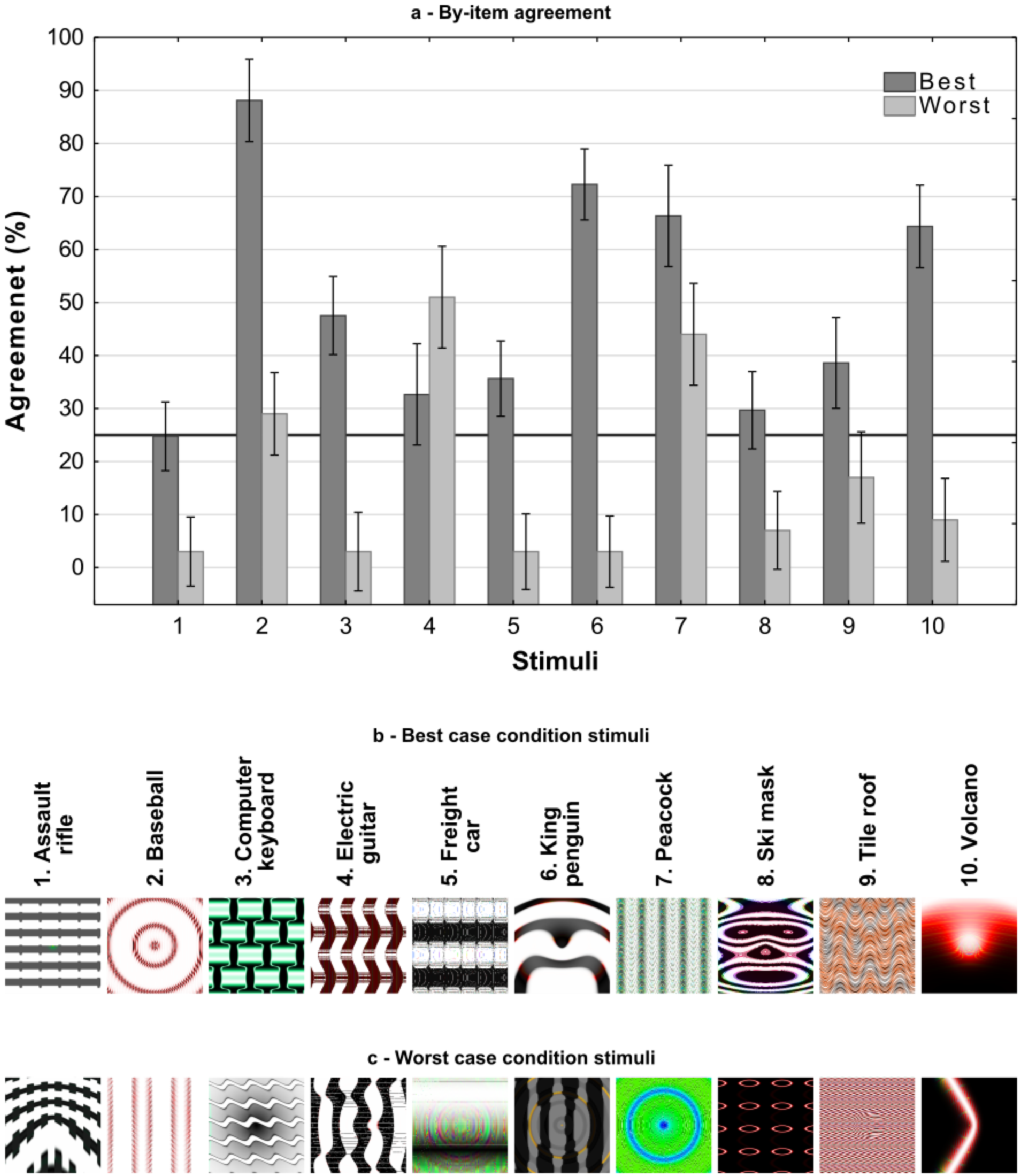


**Supplementary Figure 2.2: An item-wise breakdown of agreement levels in Experiment 2 as a function of experimental condition and category.** Average agreement levels for each category in each condition with 95% CI are presented in (a) with the black line referring to chance agreement. The best case stimuli are presented in (b), these stimuli were judged as containing the most features in common with the target category (out of 5 generated by Nguyen et al., 2015). The worst case stimuli are presented in (c), these were judged to contain the least number of features in common with the target category.

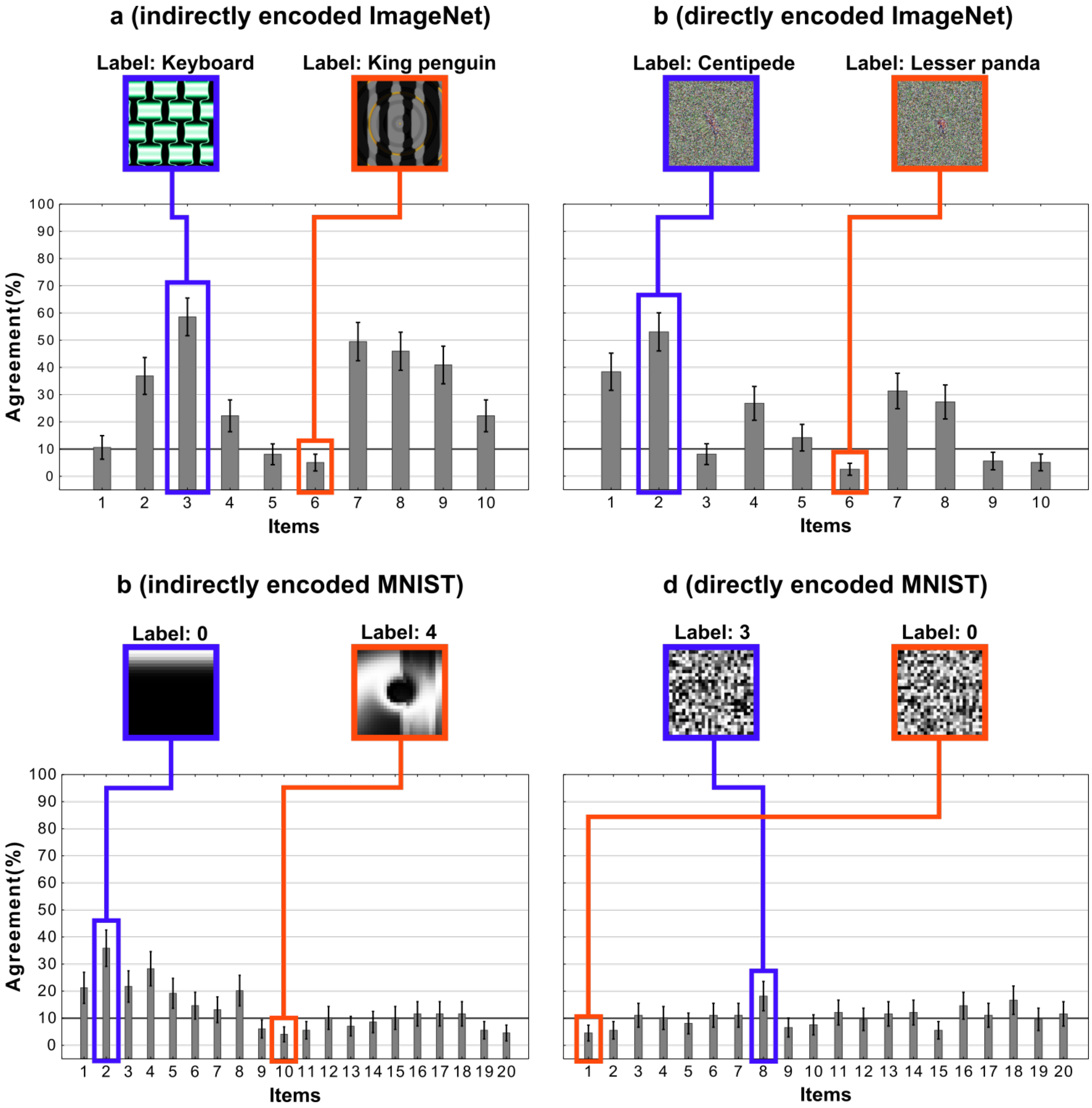


**Supplementary Figure 2.3: An item-wise breakdown of agreement levels for the four conditions in Experiment 3.** Each bar shows the agreement level for a particular image, i.e., the percentage of participants that agreed with DCNN classification for that image. Each sub-figure also shows the images that correspond to the highest (blue) and lowest (red) levels of agreement under that condition.
